# Characterization of cross-family conserved fragments

**DOI:** 10.64898/2026.09.28.754887

**Authors:** Xiang Chen, Pu Tian

## Abstract

In a companion study, we introduced ProDive and used it to identify 318,289 high-confidence cross-family segment correspondences across all 25,545 Pfam families. Here we characterize the pervasiveness and biophysical context of the resulting mapped fragment resource. Graph-based community analysis shows that segment reuse is broadly distributed across the protein family universe, with thousands of mostly small modules rather than concentration in a small number of family pairs. Comparison between Pfam families and de novo designed proteins reveals an approximately 2.1-fold enrichment of cross-family segment correspondences relative to the Pfam–Pfam background. Mapped fragments are strongly enriched in helix-dominant secondary structures with intermediate solvent exposure, occupy a narrowed sequence-context entropy regime, and show a significant shift toward lower fragment-associated contact order relative to random control windows, directly supporting a folding-compatible local structural regime. Matched annotation analyses show depletion from known domain-based interaction interfaces and mild depletion from strict DisProt disorder. Direct Start2Fold/HDX and *φ* -value comparisons are limited in scale but connect the same fragment class to folding-related protection: stability annotations support local structural stabilization that may facilitate folding, whereas folding annotations and transition-state enrichments provide more specific evidence for folding initiation or pathway involvement. Together, these results support a physical-constraint interpretation of local structural conservation, in which reusable compact fragments contribute prominently to local stability and folding-relevant local structure, with folding initiation supported in specific subsets rather than as an exclusive mechanism.

## 1 Introduction

Short local similarities across otherwise dissimilar proteins have been reported in structural and profile-based investigations of reusable segments and fragment vocabularies. ^1–6^ These observations have been discussed in the context of fragment reuse and the assembly of natural proteins from smaller structural units. ^7;8^ Yet most comparative methods in structural bioinformatics are designed to resolve similarity at the domain or fold scale, not to enumerate short local signals across otherwise unrelated families. ^9–16^ As a result, the large-scale properties of cross-family conserved fragments remain poorly characterized. In the companion ProDive study, we addressed the long-standing detection problem and generated a structurally validated Pfam-wide resource of 318,289 high-confidence local segment correspondences across all 25,545 Pfam families. Here we characterize their biophysical properties.

The present study asks a different question: what kind of molecular constraints are reflected by these fragments? The direct observation of compact local structural conservation is already informative, because it shows that short segments can recur as similar local structural solutions across different global protein contexts. That observation, however, does not by itself identify a single mechanism. Local structural conservation could arise from deep evolutionary inheritance below the domain level, convergent solutions to common physical constraints, thermodynamic stabilization, folding initiation or facilitation, or specific biochemical motifs. We therefore examine the mapped fragment resource through themed analyses of general pervasiveness, intrinsic local structural context and solvent exposure, protein-interaction-interface involvement, disorder/ordering context, and limited direct folding/stability evidence. We interpret folding relevance broadly: compact structural conservation, secondary-structure and solvent-exposure bias, sequence-context restriction, and contact order are direct biophysical evidence for locally folding-compatible fragments, whereas Start2Fold/HDX and *φ* -value measurements provide smaller but more explicit folding/stability annotations. Across these analyses, cross-family conserved fragments emerge as broadly distributed, helix-biased local features that recur frequently in de novo proteins and are depleted at known protein–protein interaction interfaces. Together, these findings support a physical-constraint interpretation in which local stability and folding facilitation stand out as major biophysical roles, while additional contributions from deep homology, convergence, and specific functional motifs remain possible. Consistent with the companion study, *segment* denotes a contiguous stretch of state positions on a pHMM, whereas *fragment* denotes the realized residue stretch on a concrete protein molecule after projection. Except where noted for graph analysis, downstream characterization in this manuscript is performed on individual projected fragments rather than fragment pairs.

## 2 Results

The Results are organized around four analytical themes. We first evaluate whether the fragment signal is broadly pervasive across natural and designed protein contexts or instead concentrated in a few function-specific family groups. We then explore the intrinsic local structural and sequence profile of the mapped fragments to identify the biophysical regime they occupy. We next examine two additional biological contexts: involvement in known protein–protein/domain interfaces and bias relative to annotated disorder. Because the preceding pervasiveness and local biophysical characterizations make folding/stability one of the most plausible general physical constraints, the final analyses explicitly ask whether the resource has direct overlap with folding- and stability-relevant annotations.

### 2.1 General pervasiveness of the fragment signal

### 2.1.1 Cross-family segment correspondences organize into pervasive graph communities

We first asked whether the Pfam-wide segment-correspondence resource is dominated by similarities confined in a relatively small number of isolated families, or instead forms a broadly distributed network of distinct recurring local similarities. We first constructed five nested score-ranked subsets of the final ProDive correspondence set (top 5%, 10%, 20%, 50%, and 100%). Each subset was then represented as a binned segment graph (Methods), where nodes are family-specific local segments and edges are retained cross-family correspondences within that subset. Because nodes arise only from correspondences included in the analyzed subset, isolated segments are not instantiated and the smallest possible reported module size is 2. Leiden community detection ^17;18^ applied independently to these subsets recovered 2,936, 4,735, 6,904, 9,654, and 11,372 modules from the top 5% subset through the full set, respectively. The largest 100 modules for the full set are shown in Fig. 1. In the full set, most reported modules remained small: 6,056 had size 2 and 1,015 had size 3, together accounting for 62.2% of all reported modules. This pattern argues against a picture in which the ProDive signal is driven by a few universal motifs or a handful of closely related clans. Instead, segment reuse appears pervasively distributed across many family contexts, with many small local modules and a limited number of larger communities. Such pervasiveness argues against explanations that reduce the resource to a few specific motif classes and motivates a broader view in which many short segments are repeatedly favored across distinct family contexts.

**Figure 1.**
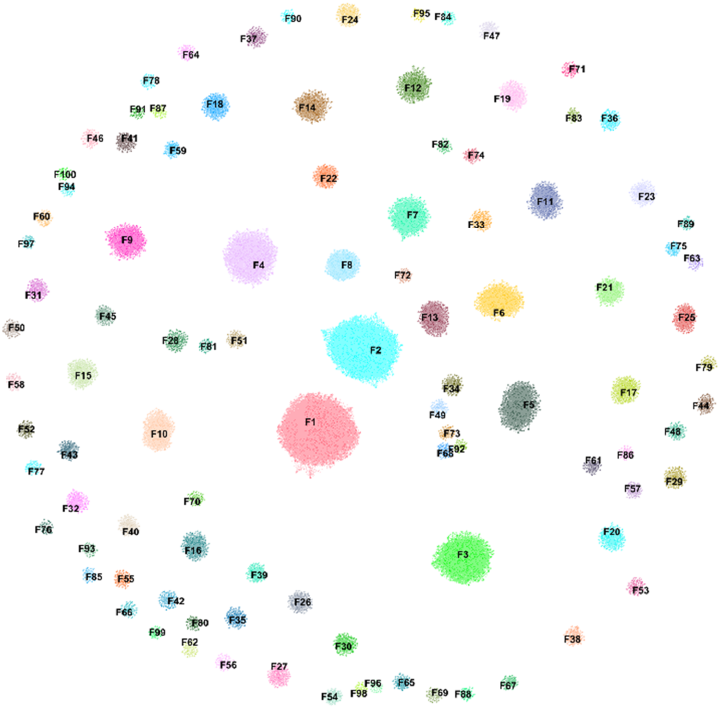
Graph-community overview of the cross-family fragment resource. Shown is the Gephi visualization of the 100 largest Leiden communities in the full fragment graph. Each node represents a binned, family-specific segment; node weights summarize how many overlapping family mappings collapsed onto that segment node, and edges represent identified cross-family correspondences. For visual clarity, only intra-community connections are shown. The figure illustrates that the resource is distributed across many modules rather than dominated by a few giant components.

#### 2.1.2 Enrichment in de novo designed proteins

De novo designed proteins provide a test of whether cross-family conserved fragments behave as reusable local structural units in designable protein space. Trained design procedures may naturally learn cross-family conserved fragments because each such fragment appears in multiple family contexts, making it available as a repeated local sequence–structure pattern even when training sets are deduplicated by whole sequence or whole-domain similarity. Family-specific positions, in contrast, should contribute less consistently to the learned design prior. We therefore asked whether the ProDive fragment class is over-represented in de novo designed proteins relative to the Pfam–Pfam background.

We collected 1,927 RCSB entries annotated as de novo designed proteins, yielding 2,249 chains after chain splitting. ^19^ Running these chains against the Pfam library with the same ProDive workflow produced approximately 117,875 de novo–Pfam diagonal paths involving 1,773 chains and 9,822 Pfam families. The comparison-normalized path rate was approximately 2.05 *×* 10^*−*3^ for de novo–Pfam versus 9.76 *×* 10^*−*4^ for the unordered Pfam–Pfam background, an enrichment of about 2.1-fold. This enrichment is not explained by a trivial relaxation of structural conservation. Of 101,864 de novo–Pfam pHMM segment correspondences submitted for structural projection, 72,547 passed structure availability, quality, coordinate-mapping, and coverage filters and yielded representative-chain fragment alignments. Within this structurally evaluable subset, 63,529 alignments occupied the short compact-core band evident in the structural-validation map, corresponding to 8–13 aligned residues in Fig. 2. Length-stratified random controls were significant across these core lengths (all *p <* 10^*−*170^; Supplementary Section S3).

**Figure 2.**
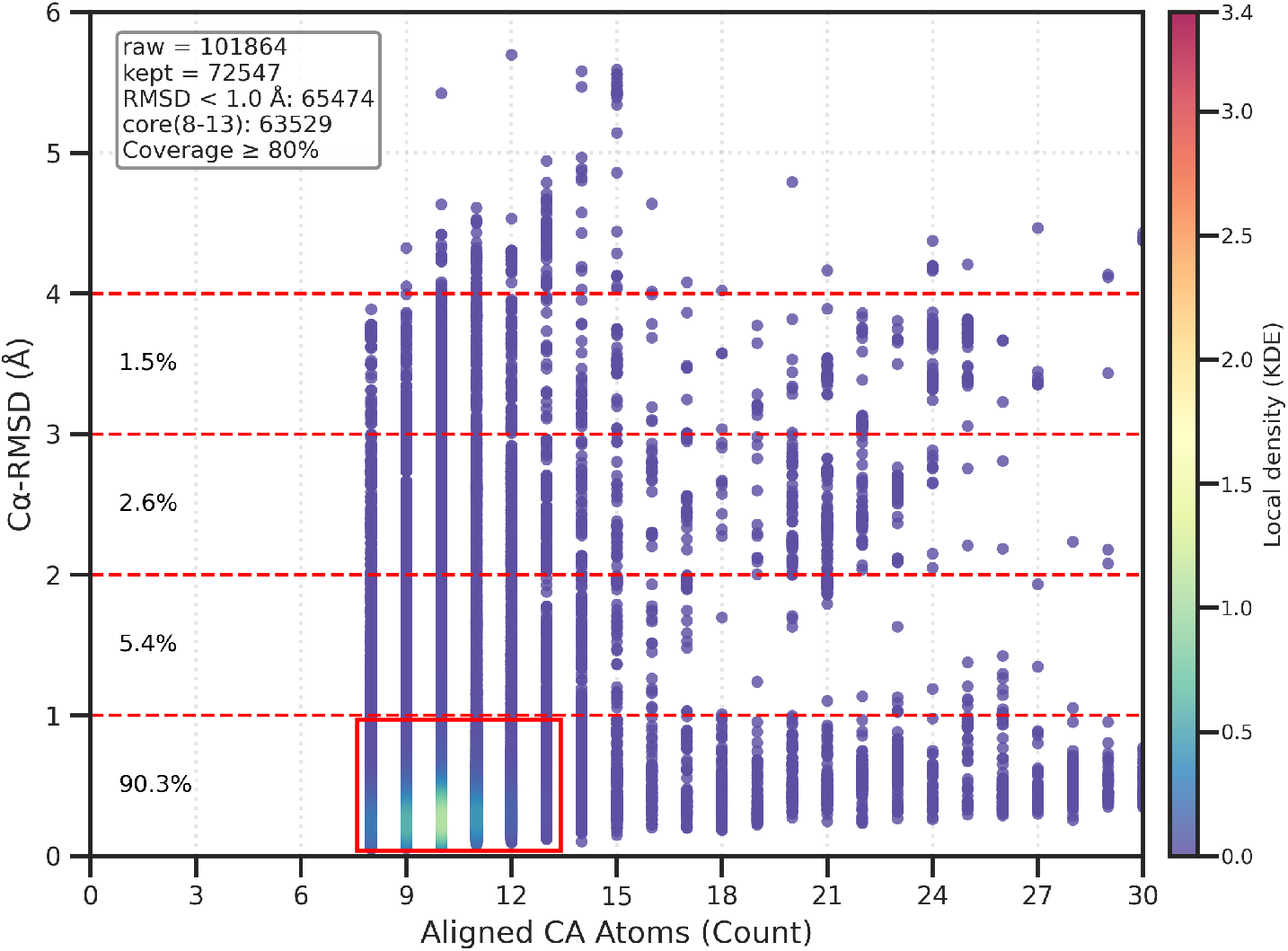
Structural validation of de novo–Pfam fragment correspondences. The distribution of validated fragment pairs is shown as aligned C_*α*_ atom count versus C_*α*_ RMSD. The red box highlights the short compact-core band evident in the validation distribution, corresponding to 8–13 aligned residues. Percentage of RMSD in (0-1], (1-2], (2-3] and (3-4] are shown in the figure. The concentration of de novo hits in this regime and the highly significant length-stratified random controls show that the de novo enrichment is not a low-specificity by-product of the screening procedure.

Designed proteins are not expected to be ordinary whole-domain homologs of the Pfam families. Nevertheless, the de novo–Pfam scan recovered the same compact local fragment correspondences at an elevated comparison-normalized rate. This supports subdomain-scale reuse and learnability: ProDive fragments appear to capture reusable local sequence–structure patterns that can be repeatedly available to design procedures, especially when those procedures are trained on or informed by natural structural data. The result should not be read as independent rediscovery from first principles or deliberate reuse of natural fragments; rather, it shows that whole-sequence or whole-domain deduplication is insufficient to remove subdomain-scale fragment reuse. Together, the graph and de novo analyses show that the fragment signal is broadly distributed across Pfam family space and over-represented in designable protein space.

### 2.2 Intrinsic local structural and biophysical profile

#### 2.2.1 Helix-dominance with intermediate solvent exposure

After graph community analysis showed broad participation across Pfam family space and the de novo comparison showed enrichment in designable protein space, we next asked what local physical environment the mapped fragments occupy. In the Pfam–Pfam set, 79.43% of observed ProDive fragments were helix-dominant compared with 46.36% in same-chain, length-matched random controls. In the de novo– Pfam comparison, similar bias was observed (Supplementary Fig. S1A,C): 85.51% of observed fragments were helix-dominant versus 71.51% for matched random windows. Sheet-dominant, coil/loop-dominant, and mixed classes were correspondingly depleted in both analyses (Fig. 3A,C; Supplementary Fig. S1A,C).

**Figure 3.**
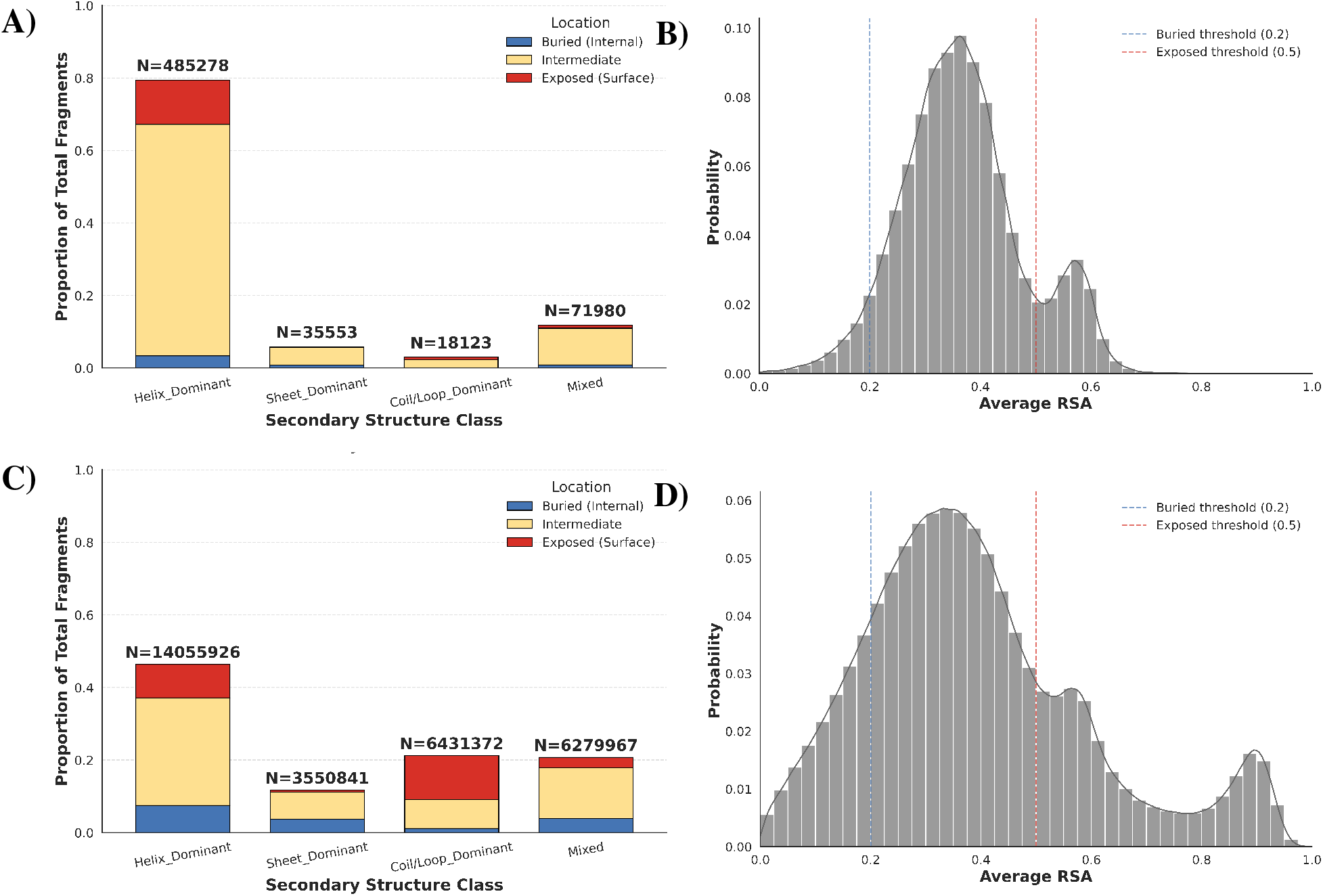
Structural context of Pfam–Pfam fragment correspondences. (A,B) Secondary-structure composition and mean RSA distribution for observed ProDive fragments. (C,D) Matched random controls drawn from the same structure chains. Observed ProDive fragments are strongly enriched in helix-dominant segments and favor the intermediate exposure regime.

The relative solvent accessibility (RSA) analysis placed mapped fragments in a middle-exposure regime. In the Pfam–Pfam set, ProDive fragments were depleted from both highly exposed and deeply buried windows and enriched at intermediate RSA relative to same-chain, length-matched controls (Fig. 3B,D). The same pattern appeared in the de novo–Pfam comparison (Supplementary Fig. S1B,D). This exposure profile is consistent with locally stable secondary-structure elements that retain partial tertiary packing, rather than fully exposed loop-like segments or uniformly buried cores.

On their own, helix enrichment and intermediate RSA do not identify a mechanism. They do, however, argue against a model in which most fragment hits reflect arbitrary flexible sequence motifs. Whatever drives cross-family fragment recurrence appears to favor compact local structures in a moderately exposed environment.

#### 2.2.2 Constrained sequence-context regime

We then asked where fragment positions fall along a sequence-context entropy spectrum: in highly predictable low-entropy contexts, in less predictable high-entropy contexts, or in an intermediate regime. Using residue-level ESM2 masked-token entropy mapped back to Pfam seed MSA columns, ^20^ we compared fragment positions with non-fragment background positions after the shared residue-to-MSA mapping and quality filters described in Methods. Detailed mapping filters and optional MSA-entropy sensitivity summaries are provided in Supplementary Section S5 and Supplementary Tables S2–S3.

The clearest signal was distributional rather than a shift to the absolute lowest entropy positions. We summarized the ESM2 distribution with three exhaustive ranges: low entropy (*≤* 1.0), intermediate entropy (1.0–2.5), and the high-entropy right tail (*>* 2.5). Fragment positions were depleted from the lowest bin (24.87% vs. 30.77% for background) and from the high-entropy right tail (16.62% vs. 22.39%), while enriched in the intermediate range (58.51% vs. 46.84%).

Thus, cross-family conserved fragments are not concentrated at either extreme of the ESM2 sequence-context entropy distribution. Instead, they occupy a narrowed regime centered on moderately constrained local contexts. This is consistent with a class of positions that is neither freely variable nor maximally predictable from local sequence context.

### 2.3 Interaction-interface and disorder context

#### 2.3.1 Depletion from known domain-based interfaces

We next asked whether mapped fragments preferentially occur in known protein–protein or domain– domain interaction interfaces using 3did domain-interaction annotations. ^21^ Counting the two mapped fragment sides of each Pfam–Pfam correspondence separately, 56,450 of 636,578 total mapped fragments (8.87%) could be evaluated against both a 3did interaction instance and a usable experimental structure. This limited evaluable fraction mainly reflects 3did and experimental-structure coverage, so non-evaluable fragments should not be interpreted as non-interface fragments.

For each evaluable fragment, we projected the fragment residues onto the matched experimental structure and measured interface overlap as the fraction of those residues that coincided with 3did contact residues. In the thresholded analysis, we first used 30% overlap as the cutoff for interface-related fragments. Among the 56,449 fragments with valid matched controls, fewer observed fragments reached this cutoff than expected from matched random windows (22.59% vs. 34.27%; observed/random = 0.659). The corresponding non-overlapping categories showed the same shift: NonInterface fragments (*<* 30% overlap) accounted for 43,696 observed cases (77.41%) versus 65.73% expected from random windows, InterfacePartial fragments (30% to less than 50% overlap) accounted for 6,612 cases (11.71%) versus 14.64% expected, and InterfaceMajor fragments (*≥* 50% overlap) accounted for 6,141 cases (10.88%) versus 19.62% expected (Supplementary Table S6). Similar depletion held across cumulative thresholds from 10% to 50% overlap (Supplementary Table S5). Without imposing a threshold, the mean residue-level interface fraction was also lower for observed fragments than for random windows (0.1407 vs. 0.2154; paired difference = -0.0747, 95% CI -0.0763 to -0.0731; Supplementary Table S4).

The same qualitative result held across 3did contact contexts. In the inclusive subset containing inter-chain different-Pfam evidence, observed fragments again showed lower interface overlap than matched random windows (0.1841 vs. 0.3066). The strict interchain different-Pfam-only, interchain same-Pfam-only, and intrachain domain-contact-only subsets showed the same trend (Supplementary Table S7). Thus, in the experimentally evaluable 3did subset, mapped fragments were significantly depleted from known domain-based interfaces relative to matched local background. Known protein–protein/domain interfaces therefore cannot be the primary explanation for the ProDive fragment signal, although individual fragments may still function at interfaces.

#### 2.3.2 Bias relative to annotated disorder

We next examined how fragment positions relate to annotated disorder and broader disorder-associated states. Using 7,120 DisProt annotation rows with explicit residue coordinates, ^22^ we retained 865 regions that could be anchored to the Pfam seed-coordinate framework. Independently, after projecting the global ProDive resource onto the same DisProt-mappable proteins and Pfam-covered sequence space, 19,926 fragment instances were testable for residue-level overlap. This is the pre-overlap fragment test set, not the number of fragments assigned to DisProt regions: 3,605 fragment instances overlapped at least one anchored DisProt annotation, covering 352 of the 865 anchored regions (Supplementary Table S8).

Raw overlap alone was not decisive. In the strict `term_name = disorder` subset, 2,570 fragment instances overlapped anchored disorder annotations, covering 248 anchored regions. We therefore used the protein-matched, Pfam-space-matched, length-matched randomization described in Methods. Under this control, strict disorder showed mild but statistically significant depletion. The observed residue-level overlap fraction was 20.03%, whereas the random mean was 20.58% (O/E = 0.974, *z* = *−*2.75, empirical depletion *p* = 0.00599; Fig. 4). When all DisProt terms were pooled, the observed overlap was 20.80% and the random mean 21.08% (O/E = 0.987, *z* = *−*1.41, depletion *p* = 0.0829), which is close to random expectation (Supplementary Table S9).

**Figure 4.**
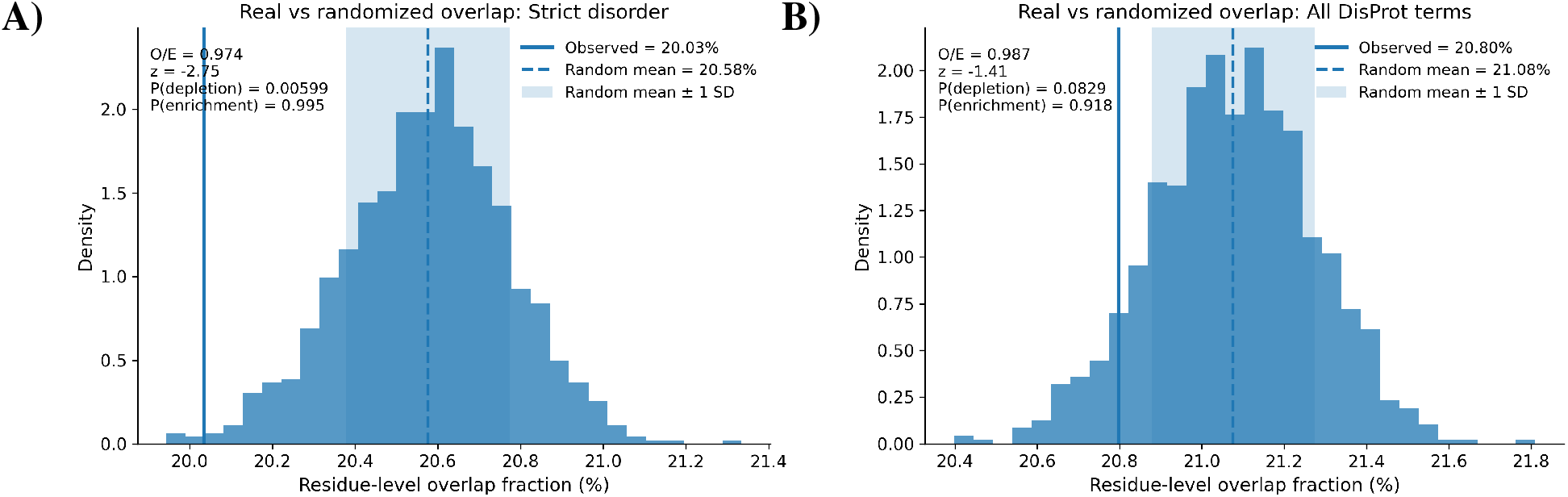
Matched-random DisProt analysis. (A) Strict-disorder overlap compared with protein-matched, Pfam-space-matched, length-matched random fragments. (B) The same comparison after pooling all DisProt terms. Observed fragments show a mild but significant depletion from strict disorder; when all DisProt terms are pooled, the overlap is close to random expectation.

The current DisProt evidence therefore indicates a mild bias away from canonical disorder, while overlap with the broader DisProt annotation universe is close to random expectation. This pattern is compatible with a local-structure interpretation without implying strong enrichment in disorder-associated states.

### 2.4 Folding relevance and local stabilization

#### 2.4.1 Fragment-associated contact order favors local-contact structure

Contact order provides a topology-based test of whether mapped fragments are preferentially coupled to long-range structural contacts or instead occupy a more local-contact regime. ^23^ Because lower contact order is linked to simpler local contact formation during folding, this analysis directly informs folding relevance, although it is not exclusive evidence for folding initiation. Using same-chain, same-length random windows as background and the formal definitions in Supplementary Section S8, observed fragments showed a small but consistent shift toward lower fragment-associated relative contact order (rFCO; Fig. 5). At an 8 Å contact cutoff, the observed rFCO mean was 0.1369 compared with 0.1436 for matched random windows, and the observed median was 0.1130 compared with 0.1376. At 6 Å, the corresponding means were 0.1248 and 0.1320. The complementary long-range contact fraction, computed from the same fragment-associated contact set, was also slightly lower for observed fragments than for matched random windows (0.5837 vs. 0.5889 at 8 Å; 0.5369 vs. 0.5435 at 6 Å; Supplementary Table S10).

**Figure 5.**
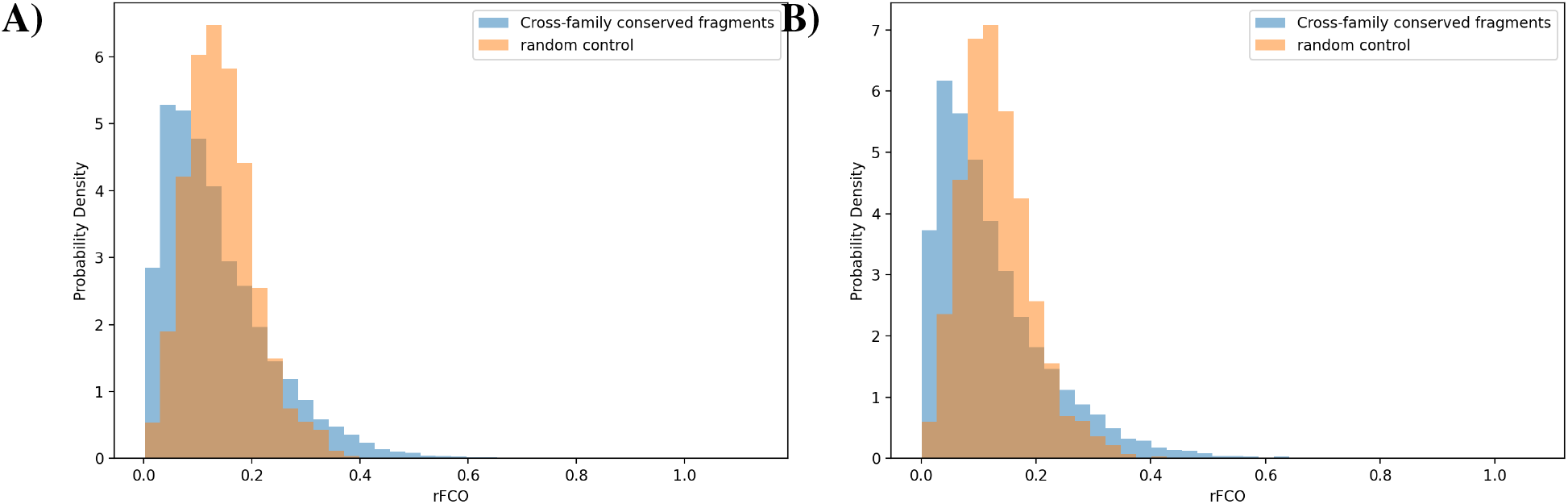
Fragment-associated contact-order distributions. (A) rFCO distributions using an 8 Å contact cutoff. (B) rFCO distributions using a 6 Å contact cutoff. Cross-family conserved fragments show a modest left shift relative to same-chain, same-length random controls under both contact definitions, while retaining a small high-rFCO right tail.

Thus, the bulk fragment population is mildly biased toward local-contact structure rather than long-range topological anchoring. A small high-rFCO tail remained, indicating that a minority of fragments may participate in long-range packing or anchoring roles. These high-rFCO fragments are less naturally interpreted as local folding initiators, because their structural context depends on contacts to distant sequence regions. Instead, they likely represent conserved local elements whose role is coupled to tertiary packing, native-state architecture, or later-stage structural consolidation. This mixed pattern is consistent with the broader structural results: most cross-family conserved fragments behave as compact local elements with direct relevance to folding and local stability, while a smaller subset is coupled to larger topological organization.

#### 2.4.2 Start2Fold/HDX annotations support local stability with folding-associated subsets

We next compared mapped fragments with Start2Fold annotations derived from HDX experiments, which provide residue-level measurements related to folding, stability, and exchange protection. ^24;25^ The Start2Fold labels are best interpreted by experimental context rather than as mutually exclusive biological roles. Folding annotations report protection acquired during refolding and therefore provide the more direct evidence for folding initiation. Stability annotations report protection in native or partially folded states; they support local structural stabilization that may facilitate folding, while also allowing roles in native-state stability beyond folding facilitation.

This analysis was necessarily small: 222 fragments could be projected into Start2Fold-associated sequence space. Nevertheless, a randomization matched by protein, seed interval, and fragment length showed higher overlap for observed fragments than expected by chance. The mean overlap fraction with any annotation was 0.398 for observed fragments versus 0.345 for random windows; fragments with at least two annotated residues occurred in 82.0% of observed cases versus 72.0% of random windows (Supplementary Table S11).

The class distribution showed that stability-context protection was the prevalent Start2Fold/HDX signal. Among all mapped fragments, 42.3% overlapped only stability-context annotations and another 38.3% overlapped both stability- and folding-context annotations, so most annotated fragments included a stability-context component. Folding-context-only cases were rare (0.45%). The dominant class was `Stab_MEDIUM` (62.6% observed versus 35.3% random). At the same time, folding-associated combinations were not absent: the combination of `Fold_EARLY`, `Fold_INTER`, and `Stab_MEDIUM` occurred in 7.66% of observed fragments but only 0.54% of random windows (Supplementary Table S12). These data support local stability/protection as the dominant direct HDX signal, with folding initiation or pathway involvement appearing in specific annotated subsets. The small number of Start2Fold-mappable proteins and the concentration of hits in a few proteins make this a direct but limited validation.

#### 2.4.3 Limited transition-state concordance

To obtain an additional direct link to folding-relevant positions, we compared mapped fragments with published residue-level *φ* -value datasets. Candidate datasets were identified from the Protein Folding Database and primary source studies. ^26–30^ Four proteins could be aligned cleanly to the Pfam seed framework: I*κ*B*α* (PF00023), NBR1-PB1 (PF00564), LysM (PF01476), and PSBD/POB* (PF02817). Using the matched randomization and high-*φ* density statistic described in Methods, observed fragment positions were compared with same-protein, same-domain random fragments (Supplementary Table S13). The result is heterogeneous but informative. I*κ*B*α* showed strong positive enrichment relative to random (observed 0.145 vs. random 0.084, *p* = 0.0001), as did NBR1-PB1 (0.260 vs. 0.116, *p* = 0.0001). LysM was slightly above random but not significant (0.359 vs. 0.328, *p* = 0.172). PSBD showed the opposite pattern, with observed fragments below random expectation (0.016 vs. 0.095, *p* = 1.000).

The evidence should therefore be separated by scale. Compact structural conservation, helix enrichment, intermediate RSA, narrowed sequence-context entropy, and the population-level contact-order shift support a folding-compatible local structural regime. Start2Fold/HDX and *φ* -value analyses are more limited: they provide direct experimental annotation tests only for the small set of proteins in which residue-level protection or folding-transition data could be mapped. Within that restricted set, Start2Fold/HDX links some fragments to protected or stabilized local structure, and *φ* -values show transition-state concordance in I*κ*B*α* and NBR1-PB1 but not uniformly across all systems. The high-rFCO tail also cautions against treating all fragments as local folding initiators, because some conserved fragments participate in long-range packing contexts. These results support a model in which compact conserved local structures often contribute to local stability and folding-compatible structure, with folding initiation or pathway involvement supported in specific subsets rather than as a universal mechanism.

## 3 Discussion

The main implication of this study is that cross-family conserved fragments represent a pervasive layer of protein structural organization below the domain scale. Their conservation is not adequately explained by known interaction interfaces, annotated disorder, or a small set of recurrent motifs. Instead, the combined evidence supports a physical-constraint model in which short compact local structures are repeatedly favored across otherwise distinct family contexts.

Within this model, local stabilization and folding facilitation are closely linked but not identical. Mapped fragments are compact, structurally conserved, helix-biased, moderately exposed, sequence-constrained, and mildly shifted toward lower fragment-associated contact order. This profile is consistent with locally stable, folding-compatible structure, but it does not require every fragment to act as a folding nucleus. Start2Fold/HDX and *φ* -value comparisons add smaller but more direct evidence that specific subsets overlap protected local structure and transition-state-relevant residues. The high-contact-order tail reinforces this plural interpretation: a minority of fragments is involved in long-range packing contexts more consistent with tertiary architecture or native-state consolidation than classic local folding initiation.

The scope of this study is the Pfam-seed-pHMM-derived ProDive resource. This scope is deliberate: Pfam provides a standardized family-wide probabilistic coordinate system in which local cross-family correspondences can be detected and characterized at scale. Within this framework, structural, sequence, interface, disorder, contact-order, and folding-annotation analyses give a consistent picture of the resulting fragment resource. At the same time, family granularity, seed-alignment composition, sequence weighting, and pHMM diffusivity may influence which local signals are detectable. Extending the analysis to alternative family models, clustering schemes, or structure-defined family representations is therefore a natural next step for distinguishing model-stable fragments from fragments specific to the current Pfam representation.

The main limitation in assigning folding mechanism is the scarcity of direct experimental folding and stability annotations. The Start2Fold/HDX and *φ* -value analyses provide useful direct support, but the available overlap sets are small and concentrated in a limited number of proteins; therefore they should be read as mechanistic consistency tests rather than population-level quantitative estimates. Expanding residue-resolved HDX, folding-kinetic, and *φ* -value datasets across more Pfam families is the key future step for testing how often conserved fragments stabilize native or partially folded states, participate in folding initiation, or instead contribute to tertiary packing, deep homology, or convergent reuse in individual systems.

The de novo result has practical implications for protein design. If cross-family conserved fragments represent densely populated local optima in foldable sequence space, then design pipelines will repeatedly rediscover them, and whole-sequence deduplication will not remove this local reuse. That is not necessarily a weakness. It may instead provide a route to more interpretable design constraints and fragment-level diagnostics for evaluating whether a designed protein occupies familiar or unusual local-structure space.

## 4 Conclusion

The ProDive fragment resource reveals a pervasive subdomain layer of local structural conservation across protein-family space. Mapped fragments recur across natural Pfam families and de novo designed proteins, favor compact helix-rich environments with intermediate solvent exposure, occupy a narrowed sequence-context entropy regime, and are depleted from known domain-interface residues and strict disorder. Their contact-order profile and limited direct Start2Fold/HDX and *φ* -value overlaps support a folding-compatible local stability regime, while leaving room for deep homology, convergent reuse, tertiary packing, and specific functional motifs in individual cases. Overall, cross-family conserved fragments are best interpreted as reusable local structural solutions shaped by physical constraints, with local stability and folding-compatible structure as prominent but not exclusive contributions.

## 5 Methods

### 5.1 Summary of the ProDive fragment resource

All analyses in this study used the final clean-path set from the companion ProDive methods study: 318,289 high-confidence Pfam–Pfam local correspondences and their associated projection tables. The algorithmic derivation of this resource, the detection-parameter choices used to generate it, structure-based validation of the 318,289 correspondences, and computation-cost benchmarks for the Pfam-wide scan are reported in the companion study and its Supplementary Information; the analyses here begin from the compact filtered correspondence resource. Downstream analysis subset sizes are summarized in Supplementary Section S1 and Supplementary Table S1.

### 5.2 Pervasiveness revealed by graph community analysis

For graph analysis, the final correspondence set was sorted by completeness-adjusted score and converted into nested top 5%, 10%, 20%, 50%, and 100% subsets. Each subset was represented as a binned segment graph whose nodes are family-specific local segments and whose edges are retained cross-family correspondences. Leiden community detection ^17;18^ with the CPM objective (*γ* = 0.05) was applied independently to each subset; binning and duplicate-collapse rules are described in Supplementary Section S2.

### 5.3 De novo protein dataset and enrichment analysis

Proteins annotated as de novo designed were collected from the RCSB Protein Data Bank and split into chain-level queries. ^19^ Each chain was compared against the Pfam family library with the same ProDive workflow used to generate the input resource. Enrichment was calculated as retained paths per possible comparison in the de novo–Pfam scan relative to the Pfam–Pfam background. Structural validation used the projection, coverage, and quality filters from the companion study; denominator definitions and length-stratified controls are described in Supplementary Section S3.

### 5.4 Secondary-structure and RSA analysis

Mapped fragments were projected onto available structures and annotated by DSSP secondary structure and residue-level relative solvent accessibility (RSA). ^31^ We summarized each fragment by its majority secondary-structure class and mean RSA. Length-matched random controls were drawn from the same structure chain and generated separately for the Pfam–Pfam and de novo–Pfam analyses; sampling details are provided in Supplementary Section S4.

### 5.5 ESM2–MSA entropy analysis

Residue-level ESM2 masked-token entropy was computed with the ESM2 `esm2_t36_3B_UR50D_MLM` masked-language-model checkpoint. ^20^ For each representative sequence, each position was masked individually while all other positions were left unchanged. Entropy was calculated over the model probabilities for the 20 standard amino acids. The resulting entropies were mapped to Pfam seed MSA columns and compared between fragment and non-fragment positions using fixed ESM2 entropy bins. Mapping filters, background definition, and optional MSA-entropy sensitivity summaries are provided in Supplementary Section S5 and Supplementary Tables S2–S3.

### 5.6 3did domain-based interface localization analysis

Interface analysis used 3did contact residues grounded to seed-sequence and experimental-structure coordinates rather than external Pfam pHMM state positions. ^21^ Each Pfam–Pfam correspondence contributes two mapped fragments, one from each family. Interface fraction was defined as the fraction of residues in a mapped fragment or matched random window that overlapped 3did contact residues after projection to the same experimental structure. Matched random controls used same-length windows from the same Pfam seed-mapped range and were evaluated against the same structure and contact-residue background. Mapping filters, thresholded categories, matched-control inference, and contact-context sensitivity analyses are described in Supplementary Section S6 and Supplementary Tables S4–S7.

### 5.7 DisProt overlap and matched random controls

DisProt annotated regions with explicit residue coordinates were mapped to Pfam seed sequences by UniProt accession and residue-range overlap. ^22^ ProDive fragments were projected into the same seed-coordinate framework. Randomization preserved protein identity, Pfam-covered analyzable interval, and fragment length. Statistical significance was assessed with a one-sided Monte Carlo matched-randomization test using the union residue-level overlap fraction as the statistic. Annotation levels, empirical-tail definitions, and raw mapping counts are described in Supplementary Section S7 and Supplementary Tables S8–S9.

### 5.8 Fragment-associated contact-order analysis

Mapped fragments with usable experimental structures were projected to PDB residue coordinates and deduplicated by identical realized fragment before summary analysis. Contacts were identified using C_*β*_ atoms when available and C_*α*_ atoms for glycine, with 8 Å and 6 Å distance cutoffs. Fragment-associated relative contact order (rFCO) and long-range contact fraction were computed from contacts involving the mapped fragment and compared with same-chain, same-length random windows. Deduplication criteria, formal metric definitions, and high-rFCO tail rules are provided in Supplementary Section S8 and Supplementary Table S10.

### 5.9 Start2Fold/HDX overlap analysis

Start2Fold annotations were used as residue-level folding and stability references. ^24^ Mapped fragments from associated Pfam families were projected into the relevant seed-sequence coordinate system and compared with protein-, interval-, and length-matched random windows. Overlap was summarized by residue-level overlap fraction and thresholded annotated-residue counts. Class definitions and random-control summaries are provided in Supplementary Section S9 and Supplementary Tables S11–S12.

### 5.10 *φ* -value overlap analysis

Proteins with published residue-level *φ* -value measurements from the Protein Folding Database and primary source studies were cross-referenced against the Pfam seed sequences used in this study. ^26–30^ After mapping experimental numbering to the seed framework, observed fragments were compared with same-protein, same-domain random fragments of identical length. The primary statistic was high-*φ* position density (*φ ≥* 0.5); system-specific intervals and summaries are provided in Supplementary Section S10 and Supplementary Table S13.

## Code availability

The ProDive source code is maintained in a private GitHub repository at https://github.com/putian74/prodive/tree/main/prodive-fragment-analysis.

## Data Availability Statement

Pfam seed alignments and family models are available from the Pfam database. ^11^ Structural analyses used publicly available entries from the RCSB Protein Data Bank ^19^ and, where relevant in the companion manuscript, AlphaFold structure predictions. ^32^ DisProt annotations are available from DisProt, ^22^ and domain-based interaction records are available from 3did. ^21^ Processed fragment tables and downstream analysis summaries will be released with the code repository. Important data derived in this study is available at https://doi.org/10.5281/zenodo.22643519.

## Acknowledgments

No funding was available for this study. No assistance from individuals who are not listed as authors was used.

## References

[1] Iddo Friedberg and Adam Godzik. Fragnostic: walking through protein structure space. Nucleic Acids Research, 33(Web Server issue):W249–W251, 2005. doi: 10.1093/nar/gki363.

[2] Lukasz Jaroszewski, Zhanwen Li, S. Sri Krishna, Constantina Bakolitsa, John Wooley, Ashley M. Deacon, Ian A. Wilson, and Adam Godzik. Exploration of uncharted regions of the protein universe. PLOS Biology, 7(9):e1000205, 2009. doi: 10.1371/journal.pbio.1000205.

[3] Vikram Alva, Johannes Soding, and Andrei N. Lupas. A vocabulary of ancient peptides at the origin of folded proteins. eLife, 4:e09410, 2015. doi: 10.7554/eLife.09410.

[4] Noelia Ferruz, Francisco Lobos, Dominik Lemm, Saacnicteh Toledo-Patino, José Arcadio Farías-Rico, Steffen Schmidt, and Birte Höcker. Identification and analysis of natural building blocks for evolution-guided fragment-based protein design. Journal of Molecular Biology, 432(13):3898–3914, 2020. doi: 10.1016/j.jmb.2020.04.013.

[5] Sergey Nepomnyachiy, Nir Ben-Tal, and Rachel Kolodny. Complex evolutionary footprints revealed in an analysis of reused protein segments of diverse lengths. Proceedings of the National Academy of Sciences, 114(44):11703–11708, 2017. doi: 10.1073/pnas.1707642114.

[6] Kaiyu Qiu, Nir Ben-Tal, and Rachel Kolodny. Similar protein segments shared between domains of different evolutionary lineages. Protein Science, 31(9):e4407, 2022. doi: 10.1002/pro.4407.

[7] Richard V. Eck and Margaret O. Dayhoff. Evolution of the structure of ferredoxin based on living relics of primitive amino acid sequences. Science, 152:363–366, 1966. doi: 10.1126/science.152.3720.363.

[8] Vikram Alva and Andrei N. Lupas. From ancestral peptides to designed proteins. Current Opinion in Structural Biology, 48:103–109, 2018. doi: 10.1016/j.sbi.2017.11.006.

[9] Alexey G. Murzin, Steven E. Brenner, Tim Hubbard, and Cyrus Chothia. SCOP: a structural classification of proteins database for the investigation of sequences and structures. Journal of Molecular Biology, 247(4):536–540, 1995. doi: 10.1016/S0022-2836(05)80134-2.

[10] Christine A. Orengo, A. D. Michie, S. Jones, D. T. Jones, M. B. Swindells, and Janet M. Thornton. CATH—a hierarchic classification of protein domain structures. Structure, 5(8):1093–1108, 1997. doi: 10.1016/S0969-2126(97)00260-8.

[11] Alex Bateman, Lachlan Coin, Richard Durbin, Robert D. Finn, Volker Hollich, Sam Griffiths-Jones, Ajay Khanna, Mhairi Marshall, Simon Moxon, Erik L. L. Sonnhammer, David J. Studholme, Corin Yeats, and Sean R. Eddy. The Pfam protein families database. Nucleic Acids Research, 32:D138–D141, 2004. doi: 10.1093/nar/gkh121.

[12] Hua Cheng, R. Dustin Schaeffer, Yuxing Liao, Lisa N. Kinch, Jimin Pei, Shuoyong Shi, Bong-Hyun Kim, and Nick V. Grishin. ECOD: An evolutionary classification of protein domains. PLOS Computational Biology, 10(12):e1003926, 2014. doi: 10.1371/journal.pcbi.1003926.

[13] Liisa Holm and Päivi Rosenström. Dali server: conservation mapping in 3D. Nucleic Acids Research, 38(Web Server issue):W545–W549, 2010. doi: 10.1093/nar/gkq366.

[14] Yang Zhang and Jeffrey Skolnick. TM-align: a protein structure alignment algorithm based on TM-score. Nucleic Acids Research, 33(7):2302–2309, 2005. doi: 10.1093/nar/gki524.

[15] Michel van Kempen, Stephanie S. Kim, Charlotte Tumescheit, Milot Mirdita, Jeongjae Lee, Cameron L. M. Gilchrist, Johannes Söding, and Martin Steinegger. Fast and accurate protein structure search with Foldseek. Nature Biotechnology, 42(2):243–246, 2024. doi: 10.1038/s41587-023-01773-0.

[16] Johannes Söding. Protein homology detection by HMM–HMM comparison. Bioinformatics, 21 (7):951–960, 2005. doi: 10.1093/bioinformatics/bti125.

[17] Vincent A. Traag, Ludo Waltman, and Nees Jan van Eck. From Louvain to Leiden: guaranteeing well-connected communities. Scientific Reports, 9(1):5233, 2019. doi: 10.1038/s41598-019-41695-z.

[18] Vincent A. Traag, Paul Van Dooren, and Yurii Nesterov. Narrow scope for resolution-limit-free community detection. Physical Review E, 84(1):016114, 2011. doi: 10.1103/PhysRevE.84.016114.

[19] Stephen K. Burley, Charmi Bhikadiya, Chunxiao Bi, Sebastian Bittrich, Henry Chao, Li Chen, Paul A. Craig, Gregg V. Crichlow, Kenneth Dalenberg, Jose M. Duarte, Shuchismita Dutta, Maryam Fayazi, Zukang Feng, Justin W. Flatt, Sai Ganesan, Sutapa Ghosh, David S. Goodsell, Rachel Kramer Green, Vladimir Guranovic, Jeremy Henry, Brian P. Hudson, Igor Khokhriakov, Catherine L. Lawson, Yuhe Liang, Robert Lowe, Ezra Peisach, Irina Persikova, Dennis W. Piehl, Yana Rose, Andrej Sali, Joan Segura, Monica Sekharan, Chenghua Shao, Brinda Vallat, Maria Voigt, Ben Webb, John D. Westbrook, Shamara Whetstone, Jasmine Y. Young, Arthur Zalevsky, and Christine Zardecki. RCSB protein data bank (RCSB.org): delivery of experimentally-determined PDB structures alongside one million computed structure models of proteins from artificial intelligence/machine learning. Nucleic Acids Research, 51(D1):D488–D508, 2023. doi: 10.1093/nar/gkac1077.

[20] Zeming Lin, Halil Akin, Roshan Rao, Brian Hie, Zhongkai Zhu, Wenting Lu, Nikita Smetanin, Robert Verkuil, Ori Kabeli, Yaniv Shmueli, Allan dos Santos Costa, Maryam Fazel-Zarandi, Tom Sercu, Salvatore Candido, and Alexander Rives. Evolutionary-scale prediction of atomic-level protein structure with a language model. Science, 379(6637):1123–1130, 2023. doi: 10.1126/science.ade2574.

[21] Roberto Mosca, Arnaud Ceol, Amelie Stein, Roger Olivella, and Patrick Aloy. 3did: a catalogue of domain-based interactions of known three-dimensional structure. Nucleic Acids Research, 42(D1): D374–D379, 2014. doi: 10.1093/nar/gkt887. URL https://3did.irbbarcelona.org/.

[22] Federica Quaglia, Bálint Mészáros, Edoardo Salladini, et al. DisProt in 2022: improved quality and accessibility of protein intrinsic disorder annotation. Nucleic Acids Research, 50(D1):D480–D487, 2022. doi: 10.1093/nar/gkab1082.

[23] Kevin W. Plaxco, Kim T. Simons, and David Baker. Contact order, transition state placement and the refolding rates of single domain proteins. Journal of Molecular Biology, 277(4):985–994, 1998. doi: 10.1006/jmbi.1998.1645.

[24] Rita Pancsa, Mihaly Varadi, Peter Tompa, and Wim F. Vranken. Start2Fold: a database of hydrogen/deuterium exchange data on protein folding and stability. Nucleic Acids Research, 44(D1): D429–D434, 2016. doi: 10.1093/nar/gkv1185.

[25] Yawen Bai. Protein folding pathways studied by pulsed- and native-state hydrogen exchange. Chemical Reviews, 106(5):1757–1768, 2006. doi: 10.1021/cr040432i.

[26] Kate F. Fulton, Glyn L. Devlin, Rachel A. Jodun, Linda Silvestri, Stephen P. Bottomley, Alan R. Fersht, and Ashley M. Buckle. PFD: a database for the investigation of protein folding kinetics and stability. Nucleic Acids Research, 33(Database issue):D279–D283, 2005. doi: 10.1093/nar/gki016.

[27] Ingrid DeVries, Diego U. Ferreiro, Ignacio E. Sánchez, and Elizabeth A. Komives. Folding kinetics of the cooperatively folded subdomain of the IκBα ankyrin repeat domain. Journal of Molecular Biology, 408(1):163–176, 2011. doi: 10.1016/j.jmb.2011.02.021.

[28] Ping Chen, Clare-Louise Evans, Jonathan D. Hirst, and Mark S. Searle. Structural insights into the two sequential folding transition states of the PB1 domain of NBR1 from Φ value analysis and biased molecular dynamics simulations. Biochemistry, 50(1):125–135, 2011. doi: 10.1021/bi1016793.

[29] Adrian A. Nickson, Kate E. Stoll, and Jane Clarke. Folding of a LysM domain: entropy–enthalpy compensation in the transition state of an ideal two-state folder. Journal of Molecular Biology, 380 (3):557–569, 2008. doi: 10.1016/j.jmb.2008.05.020.

[30] Timothy D. Sharpe, Neil Ferguson, Christopher M. Johnson, and Alan R. Fersht. Conservation of transition state structure in fast folding peripheral subunit-binding domains. Journal of Molecular Biology, 383(1):224–237, 2008. doi: 10.1016/j.jmb.2008.06.081.

[31] Matthew Z. Tien, Austin G. Meyer, Dariya K. Sydykova, Stephanie J. Spielman, and Claus O. Wilke. Maximum Allowed Solvent Accessibilites of Residues in Proteins. PLOS ONE, 8(11): e80635, 2013. doi: 10.1371/journal.pone.0080635.

[32] John Jumper, Richard Evans, Alexander Pritzel, Tim Green, Michael Figurnov, Olaf Ronneberger, Kathryn Tunyasuvunakool, Russ Bates, Augustin Žídek, Anna Potapenko, Alex Bridgland, Clemens Meyer, Simon A. A. Kohl, Andrew J. Ballard, Andrew Cowie, Bernardino Romera-Paredes, Stanislav Nikolov, Rishub Jain, Jonas Adler, Trevor Back, Stig Petersen, David Reiman, Ellen Clancy, Michal Zielinski, Martin Steinegger, Michalina Pacholska, Tamas Berghammer, Sebastian Bodenstein, David Silver, Oriol Vinyals, Andrew W. Senior, Koray Kavukcuoglu, Pushmeet Kohli, and Demis Hassabis. Highly accurate protein structure prediction with AlphaFold. Nature, 596(7873):583–589, 2021. doi: 10.1038/s41586-021-03819-2.

