## supplemental info for "Characterization of cross-family conserved fragments"

### Supplementary Methods and Reference Tables

#### Section S1. Resource summary and analysis subsets

This manuscript analyzes the final 318,289 Pfam–Pfam ProDive correspondences described in the companion study. Different downstream analyses use different projected subsets because sequence, structure, interface, disorder, and $\phi$-value mappings impose distinct coordinate requirements. Table S1 summarizes the main subset sizes used throughout the manuscript. Generation-cost and memory-footprint benchmarks for the underlying ProDive resource are reported in the companion ProDive methods supplement; this supplement documents downstream biological characterization analyses.

#### Section S2. Graph construction details

Graph construction was performed separately for the top 5%, 10%, 20%, 50%, and 100% score-ranked subsets of the final ProDive segment-correspondence set. A graph node is a family-specific local segment identifier, not an exact residue stretch. Operationally, each node is defined by the target Pfam family and boundary-bin identifiers for the projected segment. This binning prevents one- or two-column boundary shifts from creating separate graph nodes for otherwise equivalent local regions.

When several correspondences mapped to the same family-specific binned segment, they were collapsed into one node and recorded as node weight. Edges were stored as unique, unweighted, undirected links between the two segment nodes connected by a ProDive correspondence. Duplicate links arising from multiple records between the same two binned segment nodes were collapsed before community detection. Leiden community detection[1;2] was run with the CPM objective ($\gamma = 0.05$) on each graph subset. The visualization in the main figure uses the 100 largest communities from the full graph and displays intra-community links only, so it should be interpreted as a community overview rather than as the complete graph.

#### Section S3. De novo dataset and enrichment details

RCSB entries annotated as de novo designed proteins were split into chain-level queries before scanning.[3] Each designed chain was converted to the same chain representation used in the companion ProDive workflow and compared against the Pfam family library with the same window length, thresholding, path-building, and rescoring settings. A de novo–Pfam path was counted only after the same retained-path criteria used for the Pfam-wide resource had been applied.

Enrichment was calculated as retained paths per possible chain–family comparison in the de novo–Pfam scan and compared with the corresponding retained-path rate in the Pfam–Pfam background. The Pfam–Pfam denominator counted each unordered family pair once, giving a corrected de novo enrichment of approximately 2.1-fold. The subset sizes used in this comparison are listed in Table S1. Structural validation of de novo hits used the same segment-coverage and structural-quality filters as in the companion ProDive methods manuscript; additional de novo structural context panels are shown in Fig. S1. For the compact 8–13-aligned-residue regime shown in the main-text de novo structural-validation figure, observed and length-matched random RMSD distributions were compared separately for each length using Welch's $t$-test. The resulting values were $p = 7.37 \times 10^{-275}$ for $\ell = 8$, $p < 1 \times 10^{-300}$ for $\ell = 9$–11, $p = 3.49 \times 10^{-254}$ for $\ell = 12$, and $p = 6.63 \times 10^{-173}$ for $\ell = 13$.

#### Section S4. Secondary-structure and RSA annotation

Mapped fragments were projected onto usable structures and annotated by DSSP secondary structure and residue-level relative solvent accessibility (RSA).[4] Each fragment was assigned to one of four secondary-structure classes according to its dominant residue type: helix-dominant, sheet-dominant, coil/loop-dominant, or mixed. Mean RSA values were used to classify fragments into buried ($< 0.2$), intermediate (0.2–0.5), and exposed ($> 0.5$) environments for summary plots.

Random controls were drawn from the same structure chain and matched to the observed fragment length. Separate control sets were generated for the Pfam–Pfam and de novo–Pfam analyses so that the structural background comparison used the same chain population as the corresponding observed fragments. The de novo structural-context panels are provided as supplementary material because the main text uses the Pfam–Pfam panels as the primary structural-context figure.

#### Section S5. ESM2 entropy mapping details

Residue-level ESM2 masked-token entropy was generated with the ESM2 `esm2_t36_3B_UR50D_MLM` masked-language-model checkpoint, loaded through `AutoTokenizer` and `AutoModelForMaskedLM`.[5] For a sequence of length $L$, the inference script created $L$ separate masked inputs. In the input for position $i$, only residue $i$ was replaced by the mask token; no other residues were masked. The model logits at the masked position were converted to probabilities by softmax, and probabilities for the 20

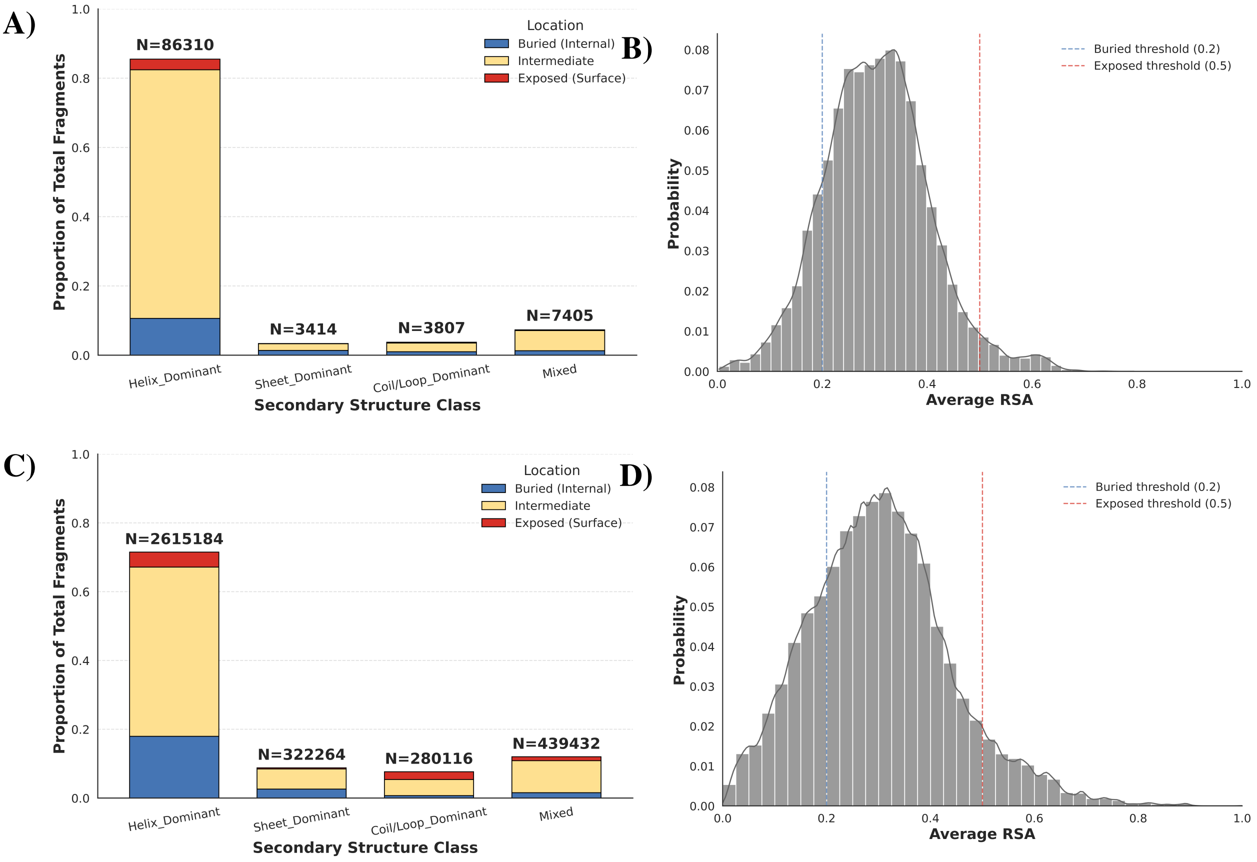


Figure S1: Structural context of de novo–Pfam fragment correspondences. (A,B) Secondary-structure composition and mean RSA distribution for observed de novo–Pfam fragments. (C,D) Matched random controls drawn from the same de novo structures. The de novo side shows the same qualitative bias toward helix-dominant fragments with intermediate exposure seen in the Pfam–Pfam analysis.

standard amino-acid tokens `ACDEFGHIKLMNPQRSTVWY` were extracted. ESM2 entropy was computed as $H_i = -\sum_a p_i(a) \log p_i(a)$ over this 20-amino-acid probability vector. Residue-level entropies were then mapped onto representative sequences and then onto Pfam seed MSA columns. Retained positions required successful residue-to-column mapping, valid ESM2 entropy, valid MSA entropy, at least five standard amino acids in the mapped column, and MSA-column gap fraction $\leq 0.80$. Per-record analyses further required at least three fragment positions and ten background positions after filtering.

The background/control set for each record consisted of non-fragment positions on the same mapped representative sequence after applying the same MSA-column and ESM2 validity filters. The main-text analysis uses the retained positions without additional MSA-entropy removal and summarizes fixed ESM2 entropy bins. For transparency, retained-position counts and ESM2-bin summaries under optional per-record MSA-entropy removal are provided as supplementary sensitivity records in Tables S2 and S3. In those sensitivity summaries, the filtering variable is Pfam seed MSA-column entropy, whereas the summarized distribution is residue-level ESM2 masked-token entropy.

#### Section S6. 3did interface mapping and random controls

3did contact records were used as residue-level interface references.[6] The counting unit was the mapped fragment: each Pfam–Pfam correspondence contributes one mapped fragment from each family. ProDive pHMM segments were projected to seed sequences and then to structure residue coordinates. Sequence-level realization had to remain within 0.8–1.2 times the nominal pHMM segment length, and mapped structural coverage had to be at least 80% of the nominal segment length.

Status categories after mapping were:

| Status | Count |
|---|---|
| `NO_COMPLEX_INSTANCE_FOR_PFAM` | 307,194 |
| `NO_MATCHING_COMPLEX_SEED_STRUCTURE` | 271,011 |
| `OK` | 56,450 |
| `COMPLEX_STRUCTURE_MAPPING_FAIL` | 1,441 |
| `LOW_STRUCTURE_COVERAGE` | 477 |
| `NO_SEED_CANDIDATE_80_120` | 5 |

Only `OK` fragments entered the interface-fraction analysis. Interface fraction was defined as the fraction of mapped residues in an observed fragment or random window that overlapped 3did contact residues. The

InterfaceMajor, InterfacePartial, and NonInterface categories are descriptive labels for observed mapped fragments, using residue-overlap thresholds of $\geq 50\%$, 30% to less than 50%, and $< 30\%$, respectively; the same thresholds were applied to matched random windows to obtain category-level background expectations. For 56,449 evaluable fragments, matched random controls sampled 200 same-length windows from the same Pfam seed-mapped range and evaluated them against the same experimental structure, contact-residue set, and multi-structure union rules. One `OK` fragment had no valid random-control set, accounting for the one-count difference between the descriptive and matched-random summaries.

Matched-control inference used continuous interface fractions and cumulative thresholds. For a threshold $t$, each observed fragment contributed an indicator for whether its interface fraction was at least $t$; its matched-random expectation was the fraction of the 200 random windows reaching the same threshold. The paired difference was averaged across fragments, and confidence intervals were calculated with clustering by ProDive Row_ID. Overall summaries are provided in Tables S4, S5, and S6.

As sensitivity checks, we separately examined 3did contact contexts, including an inclusive subset associated with at least one interchain different-Pfam contact instance, a strict interchain different-Pfam-only subset, an interchain same-Pfam-only subset, and an intrachain domain-contact-only subset. Context definitions and matched-random summaries are provided in Table S7.

#### Section S7. DisProt mapping and matched randomization

DisProt regions with explicit coordinates were mapped to Pfam seed sequences by matching UniProt accessions and testing real residue-range overlap.[7] This yielded 865 anchored analyzable regions from 7,120 original annotation rows. Independently, ProDive fragments were projected from pHMM state positions to seed-sequence residue stretches in the same proteins. The final disorder analysis used 19,926 mapped fragment instances as the pre-overlap test set. Of these, 3,605 overlapped at least one anchored DisProt annotation, covering 352 anchored regions; 2,570 overlapped the strict disorder subset, covering 248 anchored regions. These counts describe fragment instances and annotated regions separately and should not be interpreted as a one-to-one region-fragment mapping (Table S8). Randomization preserved protein identity, fragment length, and the Pfam-covered analyzable interval within each protein, and used 1,000 permutations. Residue-level overlap fractions were computed without double counting positions covered by multiple fragments.

Two annotation levels were analyzed: the strict `term_name = disorder` subset and the union of all mapped DisProt terms. Statistical significance was assessed with a one-sided Monte Carlo matched-randomization test. The test statistic was the union residue-level overlap fraction: after merging overlapping fragment-covered residues, the fraction that overlapped the tested DisProt annotation set. For depletion tests, $k$ was the number of randomizations with overlap fraction less than or equal to the observed fraction; for enrichment tests, $k$ was the number with overlap fraction greater than or equal to the observed fraction. Empirical one-sided probabilities were reported as $(k+1)/(n+1)$ with $n = 1{,}000$, which prevents zero empirical $p$-values when no matched randomization is more extreme than the observation. Compact numeric summaries are provided in Table S9.

#### Section S8. Fragment-associated contact-order analysis details

Contact-order analysis used mapped Pfam–Pfam fragments with usable experimental structures. Before summary analysis, identical realized fragments were collapsed so that repeated ProDive correspondences projecting to the same local structure did not receive repeated weight. Duplicates were defined by identical Pfam family, UniProt accession, PDB structure and chain, pHMM state start and end positions, target-sequence start and end positions, and exact mapped PDB residue-token set; the first occurrence was retained. For each mapped fragment, contacts were identified at 8 Å and 6 Å distance cutoffs, excluding near-sequence contacts within two residues. Let $F$ denote the residue set of a mapped fragment, $L$ the chain length, and $C_F$ the set of residue–residue contacts $(i, j)$ such that at least one residue lies in $F$:

$$C_F = \{(i, j) : d_{ij} \leq d_{\text{cut}},\ |i-j| > 2,\ i \in F \text{ or } j \in F\}.$$

Fragment-associated relative contact order (rFCO) was then defined as

$$\text{rFCO}(F) = \frac{1}{L|C_F|} \sum_{(i,j)\in C_F} |i-j|.$$

Thus, rFCO is the mean sequence separation of fragment-associated contacts, normalized by chain length. Long-range contact fraction is a complementary thresholded statistic computed from the same contact set:

$$\text{LRfrac}(F) = \frac{1}{|C_F|} \sum_{(i,j)\in C_F} \mathbf{1}(|i-j| \geq 12).$$

It measures the fraction of fragment-associated contacts that span at least 12 residues along the chain. rFCO and long-range contact fraction are therefore related but not identical: rFCO is a normalized continuous average, whereas long-range contact fraction is a thresholded proportion. Windows with $|C_F| = 0$ do not have defined rFCO or long-range contact fraction and were excluded from metric-specific summaries.

For each observed fragment, 1,000 candidate same-chain, same-length random windows were sampled and processed with the same contact definitions. The rFCO delta used in high-rFCO tail analyses was defined as

$$\Delta\text{rFCO}(F) = \text{rFCO}_{\text{observed}}(F) - \overline{\text{rFCO}}_{\text{matched random}}(F),$$

where the matched-random mean was calculated from valid random windows for the same fragment. Compact summaries are provided in Table S10.

High-rFCO tail subsets were defined at the 8 Å contact cutoff. The loose subset required rFCO $\geq 0.35$. The strict subset required rFCO $\geq 0.40$, rFCO delta versus the matched random mean $\geq 0.10$, empirical left-tail percentile $\geq 0.99$, and at least 10 fragment-associated contacts. The extreme subset used the same criteria but required rFCO $\geq 0.50$. These tail subsets were used to assess whether a minority of fragments showed stronger long-range topological coupling than the bulk population. Such fragments were treated as a distinct minority class, because high fragment-associated contact order is less consistent with classic local folding initiation and more consistent with tertiary packing, anchoring, or native-state architectural constraints.

#### Section S9. Start2Fold/HDX overlap analysis details

Start2Fold annotations were used as residue-level HDX-protection references.[8] Candidate fragments from Start2Fold-associated Pfam families were projected into seed-sequence coordinates, yielding 222 analyzable fragments. Random controls were matched by protein, seed sequence, analyzable seed-space interval, and fragment length, with 1,000 random windows sampled per observed fragment. Overlap was summarized as residue-level overlap fraction with any Start2Fold/HDX annotation and as thresholded hit rates requiring at least one or at least two annotated residues.

Start2Fold folding and stability labels were interpreted by experimental context. Folding labels record protection acquired during refolding and are therefore the more direct evidence for folding initiation or pathway progression. Stability labels record protection in native or partially folded states; they support local structural stabilization that may facilitate folding, but may also reflect native-state stability beyond folding facilitation. Start2Fold-derived classes were summarized as state categories, dominant class, and exact class combinations. Compact summaries are provided in Tables S11 and S12.

This direct analysis is small. Hits were concentrated in a limited number of Start2Fold proteins, especially STF0006 and STF0025, so the result should be interpreted as support for local stability/protection and folding-associated subsets in the mappable annotated set rather than as a universal statement about all fragments.

##### Section S10. $\phi$-value analysis details

The $\phi$-value randomization tested whether observed ProDive fragments were more likely than matched random fragments to cover high-$\phi$ positions within the same protein and the same Pfam seed-mapped domain interval. Candidate $\phi$-value datasets were identified from the Protein Folding Database and primary source studies.[9–13] The sampled domain intervals were 143–175 for I$\kappa$B$\alpha$ (P25963), 402–443 for LysM (P0AEZ7), 4–85 for NBR1-PB1 (Q14596), and 95–130 for PSBD/POB* (Q8ZUR6). The primary statistic was high-$\phi$ position density, defined as the number of covered positions with $\phi \geq 0.5$ divided by fragment length. For each system, random fragments were sampled within the same seed-mapped domain interval and matched to the observed fragment lengths. System-level summaries are provided in Table S13.

#### Supplementary Tables

Table S1: Main subset sizes used across downstream analyses.

| Analysis subset | Count |
|---|---:|
| Final Pfam–Pfam ProDive correspondences | 318,289 |
| Mapped fragments in Pfam–Pfam resource | 636,578 |
| 3did-evaluable fragments | 56,450 |
| 3did-evaluable fragments with valid matched random controls | 56,449 |
| Mapped fragments used in DisProt analysis | 19,926 |
| Anchored DisProt regions used in analysis | 865 |
| Contact-order deduplicated mapped fragments (8 Å) | 31,857 |
| Start2Fold/HDX analyzable mapped fragments | 222 |
| De novo designed RCSB entries | 1,927 |
| De novo chains after chain splitting | 2,249 |
| Designed chains with retained de novo–Pfam paths | 1,773 |
| Pfam families hit by de novo chains | 9,822 |

Table S2: Optional per-record MSA-entropy removal counts for the ESM2 analysis. Removal thresholds refer to the lowest Pfam seed MSA-entropy positions within each representative sequence, after residue-to-MSA mapping and quality filtering.

| MSA-low removed | Retained total | Removed total | Actual removed | Fragment retained | Background retained |
|---|---|---|---|---|---|
| 0% | 2,892,212 | 0 | 0.00% | 438,638 | 2,453,574 |
| 5% | 2,747,600 | 144,612 | 5.00% | 423,477 | 2,324,123 |
| 10% | 2,603,015 | 289,197 | 10.00% | 406,986 | 2,196,029 |
| 20% | 2,313,779 | 578,433 | 20.00% | 370,945 | 1,942,834 |
| 50% | 1,446,021 | 1,446,191 | 50.00% | 245,881 | 1,200,140 |

Table S3: Fixed ESM2 entropy-bin summaries under optional per-record MSA-entropy removal. The removal threshold is based on Pfam seed MSA entropy, whereas the three reported bins use ESM2 masked-token entropy: low ($\leq 1.0$), intermediate ($> 1.0$ and $\leq 2.5$), and high ($> 2.5$).

| Lowest MSA entropy removed | Frag low | Bg low | Frag interm. | Bg interm. | Frag high | Bg high |
|---|---|---|---|---|---|---|
| 0% | 24.87% | 30.77% | 58.51% | 46.84% | 16.62% | 22.39% |
| 5% | 23.22% | 28.50% | 59.96% | 48.71% | 16.82% | 22.79% |
| 10% | 21.58% | 26.37% | 61.37% | 50.41% | 17.05% | 23.22% |
| 20% | 18.41% | 22.48% | 63.98% | 53.38% | 17.61% | 24.14% |
| 50% | 10.83% | 13.65% | 69.24% | 58.87% | 19.93% | 27.48% |

Table S4: 3did interface summary statistics for evaluable fragments. Categorical random values are matched expectations from same-chain, same-length random windows; continuous comparisons use the residue-level interface fraction.

| Metric | Value |
|---|---|
| *Categorical overlap summary* | |
| Fragments with valid random controls | 56,449 |
| Observed NonInterface ($< 30\%$) | 43,696 (77.41%) |
| Random-window NonInterface ($< 30\%$) | 65.73% |
| Observed InterfacePartial (30% to $< 50\%$) | 6,612 (11.71%) |
| Random-window InterfacePartial (30% to $< 50\%$) | 14.64% |
| Observed InterfaceMajor ($\geq 50\%$) | 6,141 (10.88%) |
| Random-window InterfaceMajor ($\geq 50\%$) | 19.62% |
| Observed interface-related ($\geq 30\%$) | 22.59% |
| Random-window interface-related ($\geq 30\%$) | 34.27% |
| *Continuous matched-random comparison* | |
| Matched random windows | 11,289,800 |
| Observed fragment mean residue-level interface fraction | 0.1407 |
| Random-window mean residue-level interface fraction | 0.2154 |
| Paired difference (observed − random) | -0.0747 |
| 95% CI for paired difference | -0.0763 to -0.0731 |
| Directional depletion test | $p < 0.001$ |
| Observed fragment median residue-level interface fraction | 0.0000 |
| Median of per-fragment random-window mean | 0.1795 |
| Observed fragment > own random-window mean | 23.74% |

Table S5: Cumulative 3did interface-overlap thresholds with matched random controls. For each threshold, the random value is the average per-fragment probability that matched random windows reached the same residue-level interface-overlap threshold. All paired differences had depletion-direction $p < 0.001$.

| Threshold | Real count | Real | Random | Diff. | 95% CI | O/R |
|---|---|---|---|---|---|---|
| $\geq$ 10% | 21,766 | 38.56% | 50.71% | -12.15 pp | -12.52 to -11.78 pp | 0.760 |
| $\geq$ 20% | 16,894 | 29.93% | 42.23% | -12.30 pp | -12.65 to -11.95 pp | 0.709 |
| $\geq$ 30% | 12,753 | 22.59% | 34.27% | -11.68 pp | -12.00 to -11.36 pp | 0.659 |
| $\geq$ 40% | 9,293 | 16.46% | 26.46% | -10.00 pp | -10.29 to -9.72 pp | 0.622 |
| $\geq$ 50% | 6,141 | 10.88% | 19.62% | -8.74 pp | -8.99 to -8.50 pp | 0.554 |

Table S6: Non-overlapping 3did interface-overlap intervals. The table describes the distributional shift and is derived from adjacent cumulative thresholds; formal inference is based on the cumulative paired comparisons in Table S5.

| Overlap interval | Real count | Real | Random | Diff. |
|---|---|---|---|---|
| $<$ 10% | 34,683 | 61.44% | 49.29% | +12.15 pp |
| 10–$<$ 20% | 4,872 | 8.63% | 8.48% | +0.15 pp |
| 20–$<$ 30% | 4,141 | 7.34% | 7.96% | -0.62 pp |
| 30–$<$ 40% | 3,460 | 6.13% | 7.80% | -1.68 pp |
| 40–$<$ 50% | 3,152 | 5.58% | 6.84% | -1.26 pp |
| $\geq$ 50% | 6,141 | 10.88% | 19.62% | -8.74 pp |

Table S7: 3did contact-context sensitivity analysis. The inclusive interchain different-Pfam subset contains fragments with at least one interchain different-Pfam contact instance, whereas the “only” subsets contain fragments assigned exclusively to the stated context.

| Subset | N | Real mean | Random mean | Diff. | 95% CI |
|---|---|---|---|---|---|
| All evaluable fragments | 56,449 | 0.1407 | 0.2154 | -0.0747 | -0.0763 to -0.0731 |
| Any interchain different-Pfam evidence | 14,729 | 0.1841 | 0.3066 | -0.1225 | -0.1261 to -0.1189 |
| Interchain different-Pfam only | 4,152 | 0.1763 | 0.2741 | -0.0977 | -0.1037 to -0.0917 |
| Interchain same-Pfam only | 27,727 | 0.1067 | 0.1619 | -0.0552 | -0.0573 to -0.0531 |
| Intrachain domain-contact only | 10,841 | 0.1416 | 0.1985 | -0.0569 | -0.0604 to -0.0534 |

Table S8: DisProt mapping and raw overlap counts. Fragment counts are ProDive fragment instances mapped into the DisProt/Pfam coordinate space; region counts are anchored DisProt intervals.

| Quantity | Count |
|---|---|
| Original DisProt annotation rows | 7,120 |
| Anchored analyzable DisProt regions | 865 |
| Testable ProDive fragment instances | 19,926 |
| Fragment instances overlapping any DisProt term | 3,605 |
| Anchored regions covered by any fragment | 352 |
| Fragment instances overlapping strict disorder | 2,570 |
| Anchored strict-disorder regions covered by any fragment | 248 |
| Pfam-covered analyzable sequence space | 75,805 aa |
| Strict-disorder residues in Pfam-covered space | 13,004 aa |
| All-term DisProt residues in Pfam-covered space | 13,502 aa |
| Union of observed fragment-covered residues | 12,843 aa |
| Fragment-covered residues overlapping strict disorder | 2,573 aa |
| Fragment-covered residues overlapping any DisProt term | 2,671 aa |

Table S9: DisProt matched-random overlap statistics.

| Annotation level | Observed | Random mean | O/E | Depletion $p$ |
|---|---|---|---|---|
| Strict disorder | 20.03% | 20.58% | 0.974 | 0.00599 |
| All DisProt terms | 20.80% | 21.08% | 0.987 | 0.0829 |

Table S10: Fragment-associated contact-order summary. Observed fragments were compared with same-chain, same-length random windows under two contact definitions. rFCO denotes fragment-associated relative contact order.

| Metric | 8 Å | 6 Å |
|---|---|---|
| Raw input fragments | 96,092 | 96,092 |
| Fully mapped fragments | 91,294 | 91,244 |
| Deduplicated observed fragments | 31,857 | 31,836 |
| Valid random windows | 31,821,813 | 31,789,053 |
| Observed rFCO mean | 0.1369 | 0.1248 |
| Random rFCO mean | 0.1436 | 0.1320 |
| Observed rFCO median | 0.1130 | 0.0980 |
| Random rFCO median | 0.1376 | 0.1247 |
| Observed long-range contact fraction mean | 0.5837 | 0.5369 |
| Random long-range contact fraction mean | 0.5889 | 0.5435 |
| *8 Å high-rFCO tail subsets* | | |
| Loose high-rFCO subset | 1,279 | |
| Strict high-rFCO subset | 66 | |
| Extreme high-rFCO subset | 21 | |

Table S11: Start2Fold/HDX overlap summary. Random controls were matched by protein, seed-space interval, and fragment length.

| Metric | Observed | Random | Ratio |
|---|---|---|---|
| Analyzable fragments | 222 | 222,000 | – |
| Mean overlap fraction with any annotation | 0.398 | 0.345 | 1.154 |
| Fragments with $\geq 1$ annotated residue | 87.4% | 82.2% | 1.063 |
| Fragments with $\geq 2$ annotated residues | 82.0% | 72.0% | 1.139 |
| No-hit state | 18.9% | 31.7% | 0.596 |
| Single-type state | 41.9% | 32.9% | 1.274 |
| Multi-type state | 39.2% | 35.4% | 1.107 |

Table S12: Start2Fold/HDX folding- and stability-context class summaries. Class proportions are computed over the 222 observed fragments and matched random windows. Stability-context labels indicate protected native or partially folded states, not biological independence from folding.

| **Class or class combination** | **Observed** | **Random** | **Enrichment** |
|---|---|---|---|
| Stability-context-only state | 42.3% | 35.6% | 1.189 |
| Both folding and stability contexts | 38.3% | 29.9% | 1.279 |
| Folding-context-only state | 0.45% | 2.75% | 0.164 |
| `Stab_MEDIUM` dominant class | 62.6% | 35.3% | 1.776 |
| `Fold_EARLY` dominant class | 0.90% | 5.98% | 0.151 |
| `Stab_MEDIUM` exact combination | 32.4% | 22.7% | 1.429 |
| `Fold_EARLY;Fold_INTER;` `Stab_MEDIUM` exact combination | 7.66% | 0.54% | 14.167 |

Table S13: $\phi$-value randomization summary by system. High-$\phi$ density is the fraction of fragment positions with $\phi \geq 0.5$.

| **System** | **Observed density** | **Random mean** | **Right-tail *p*** |
|---|---|---|---|
| I$\kappa$B$\alpha$ | 0.145 | 0.084 | 0.0001 |
| NBR1-PB1 | 0.260 | 0.116 | 0.0001 |
| LysM | 0.359 | 0.328 | 0.172 |
| PSBD / POB* | 0.016 | 0.095 | 1.000 |

#### References

[1] Vincent A. Traag, Ludo Waltman, and Nees Jan van Eck. From Louvain to Leiden: guaranteeing well-connected communities. *Scientific Reports*, 9(1):5233, 2019. doi: 10.1038/s41598-019-41695-z.

[2] Vincent A. Traag, Paul Van Dooren, and Yurii Nesterov. Narrow scope for resolution-limit-free community detection. *Physical Review E*, 84(1):016114, 2011. doi: 10.1103/PhysRevE.84.016114.

[3] Stephen K. Burley, Charmi Bhikadiya, Chunxiao Bi, Sebastian Bittrich, Henry Chao, Li Chen, Paul A. Craig, Gregg V. Crichlow, Kenneth Dalenberg, Jose M. Duarte, Shuchismita Dutta, Maryam Fayazi, Zukang Feng, Justin W. Flatt, Sai Ganesan, Sutapa Ghosh, David S. Goodsell, Rachel Kramer Green, Vladimir Guranovic, Jeremy Henry, Brian P. Hudson, Igor Khokhriakov, Catherine L. Lawson, Yuhe Liang, Robert Lowe, Ezra Peisach, Irina Persikova, Dennis W. Piehl, Yana Rose, Andrej Sali, Joan Segura, Monica Sekharan, Chenghua Shao, Brinda Vallat, Maria Voigt, Ben Webb, John D. Westbrook, Shamara Whetstone, Jasmine Y. Young, Arthur Zalevsky, and Christine Zardecki. RCSB protein data bank (RCSB.org): delivery of experimentally-determined PDB structures alongside one million computed structure models of proteins from artificial intelligence/machine learning. *Nucleic Acids Research*, 51(D1):D488–D508, 2023. doi: 10.1093/nar/gkac1077.

[4] Matthew Z. Tien, Austin G. Meyer, Dariya K. Sydykova, Stephanie J. Spielman, and Claus O. Wilke. Maximum Allowed Solvent Accessibilites of Residues in Proteins. *PLOS ONE*, 8(11): e80635, 2013. doi: 10.1371/journal.pone.0080635.

[5] Zeming Lin, Halil Akin, Roshan Rao, Brian Hie, Zhongkai Zhu, Wenting Lu, Nikita Smetanin, Robert Verkuil, Ori Kabeli, Yaniv Shmueli, Allan dos Santos Costa, Maryam Fazel-Zarandi, Tom Sercu, Salvatore Candido, and Alexander Rives. Evolutionary-scale prediction of atomic-level protein structure with a language model. *Science*, 379(6637):1123–1130, 2023. doi: 10.1126/science.ade2574.

[6] Roberto Mosca, Arnaud Ceol, Amelie Stein, Roger Olivella, and Patrick Aloy. 3did: a catalogue of domain-based interactions of known three-dimensional structure. *Nucleic Acids Research*, 42(D1): D374–D379, 2014. doi: 10.1093/nar/gkt887. URL `https://3did.irbbarcelona.org/`.

[7] Federica Quaglia, Bálint Mészáros, Edoardo Salladini, et al. DisProt in 2022: improved quality and

accessibility of protein intrinsic disorder annotation. *Nucleic Acids Research*, 50(D1):D480–D487, 2022. doi: 10.1093/nar/gkab1082.

[8] Rita Pancsa, Mihaly Varadi, Peter Tompa, and Wim F. Vranken. Start2Fold: a database of hydrogen/deuterium exchange data on protein folding and stability. *Nucleic Acids Research*, 44(D1): D429–D434, 2016. doi: 10.1093/nar/gkv1185.

[9] Kate F. Fulton, Glyn L. Devlin, Rachel A. Jodun, Linda Silvestri, Stephen P. Bottomley, Alan R. Fersht, and Ashley M. Buckle. PFD: a database for the investigation of protein folding kinetics and stability. *Nucleic Acids Research*, 33(Database issue):D279–D283, 2005. doi: 10.1093/nar/gki016.

[10] Ingrid DeVries, Diego U. Ferreiro, Ignacio E. Sánchez, and Elizabeth A. Komives. Folding kinetics of the cooperatively folded subdomain of the I$\kappa$B$\alpha$ ankyrin repeat domain. *Journal of Molecular Biology*, 408(1):163–176, 2011. doi: 10.1016/j.jmb.2011.02.021.

[11] Ping Chen, Clare-Louise Evans, Jonathan D. Hirst, and Mark S. Searle. Structural insights into the two sequential folding transition states of the PB1 domain of NBR1 from $\Phi$ value analysis and biased molecular dynamics simulations. *Biochemistry*, 50(1):125–135, 2011. doi: 10.1021/bi1016793.

[12] Adrian A. Nickson, Kate E. Stoll, and Jane Clarke. Folding of a LysM domain: entropy–enthalpy compensation in the transition state of an ideal two-state folder. *Journal of Molecular Biology*, 380 (3):557–569, 2008. doi: 10.1016/j.jmb.2008.05.020.

[13] Timothy D. Sharpe, Neil Ferguson, Christopher M. Johnson, and Alan R. Fersht. Conservation of transition state structure in fast folding peripheral subunit-binding domains. *Journal of Molecular Biology*, 383(1):224–237, 2008. doi: 10.1016/j.jmb.2008.06.081.
